## Supplementary Material for "Causal role of the angular gyrus in insight-driven memory reconfiguration"

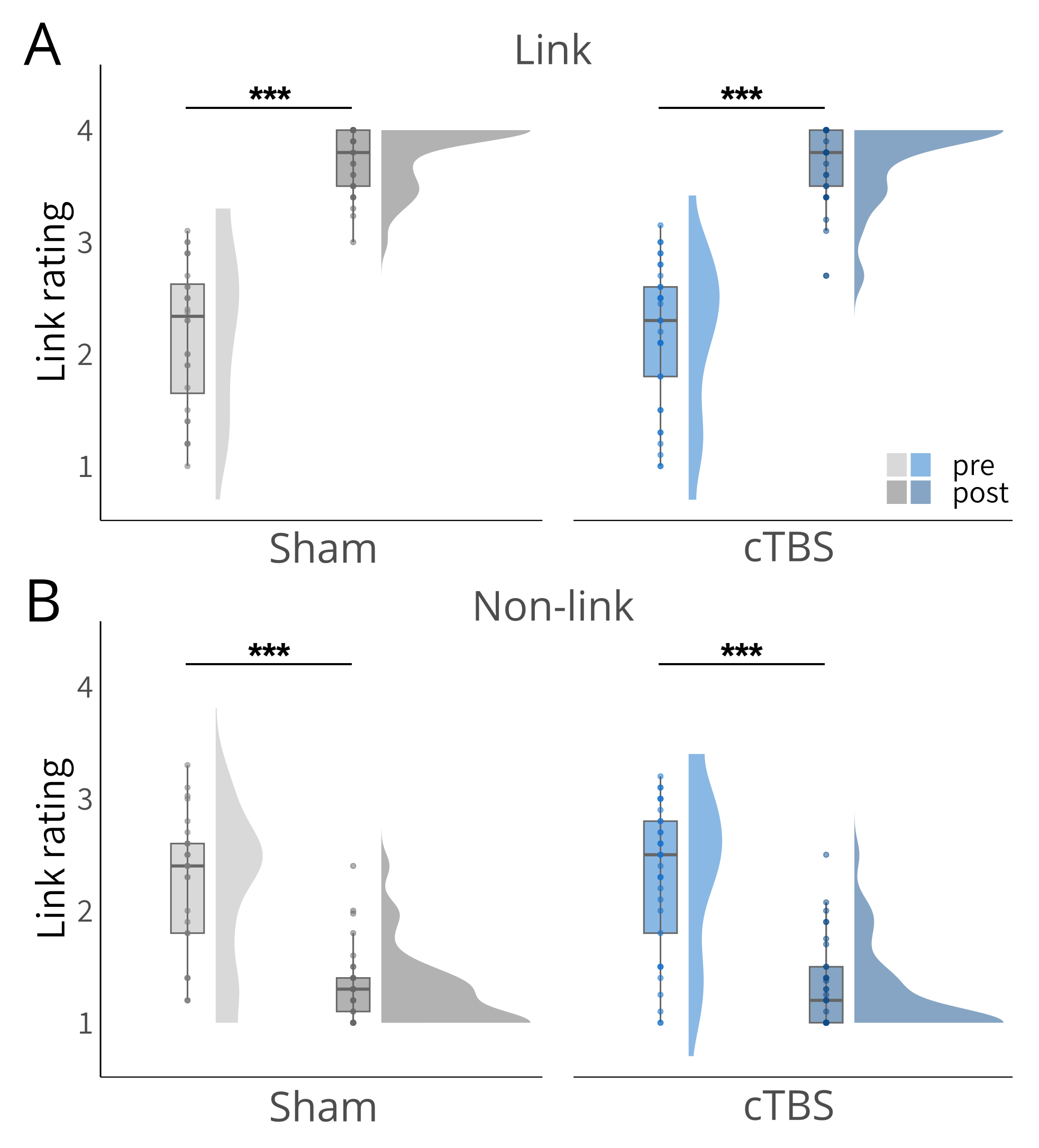

Figure 2 – figure supplement 1. Results of the narrative-insight task (NIT). **A**, Significant increases in ratings for linked events from the pre- to the post-phase for both, the sham and the cTBS groups. There were no significant group differences. **B**, Significant decreases in ratings for non-linked events from the pre- to the post-phase for both, the sham and the cTBS groups. There were no significant group differences. Boxplots show the median link ratings for each group at each time point. Boxplot whiskers extend to the minimum or maximum value within 1.5 times the interquartile range. Points within the boxplot indicate individual data points in each group. Density plots indicate data distribution per group and time. Statistical differences stem from pairwise post-hoc tests of marginal means. ****p* < .001.

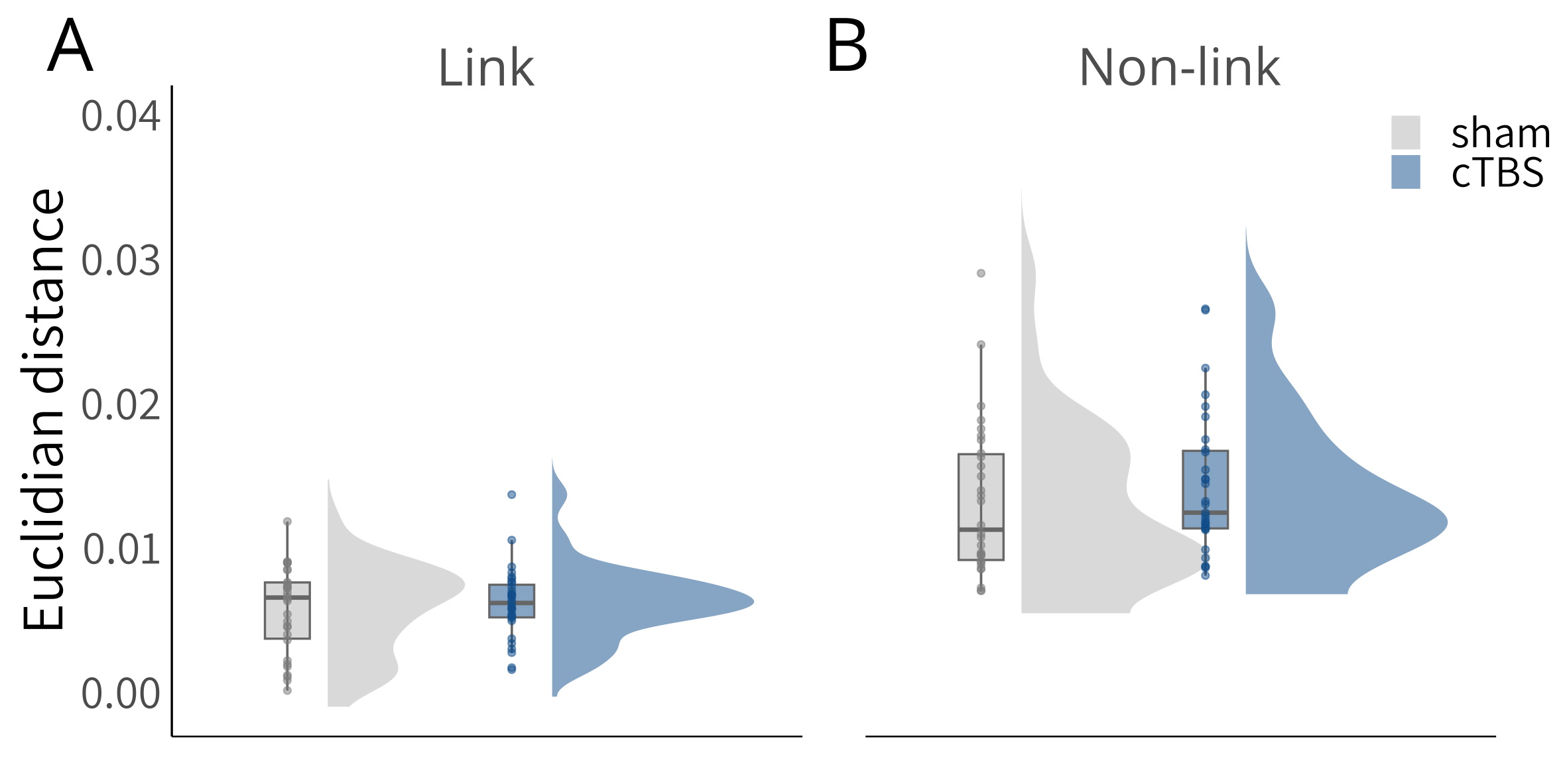

Figure 2 – figure supplement 2. Results of the multiple-arrangements task (MAT). Both groups arranged linked events closer to each other than non-linked events, without differences between groups. Boxplots show the median Euclidian distance for each group at each time point. Boxplot whiskers extend to the minimum or maximum value within 1.5 times the interquartile range. Points within the boxplot indicate individual data points in each group. Density plots indicate data distribution per group.

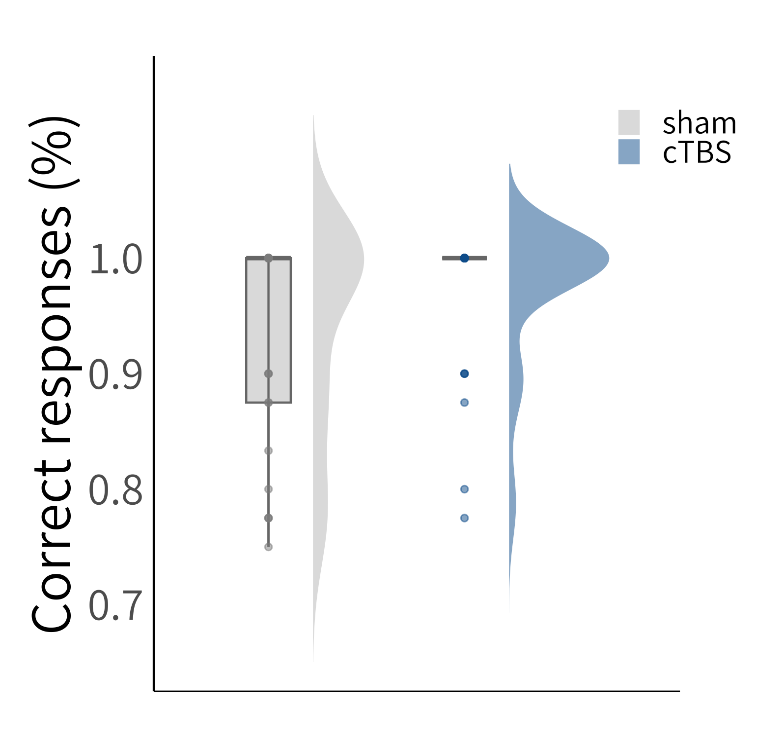

Figure 2 – figure supplement 3. Results of the forced-choice recognition task. Both groups performed very well in this task, without group differences. Boxplots show the median correct responses (%) for each group. Boxplot whiskers extend to the minimum or maximum value within 1.5 times the interquartile range. Points within the boxplot indicate individual data points in each group. Density plots indicate data distribution per group.

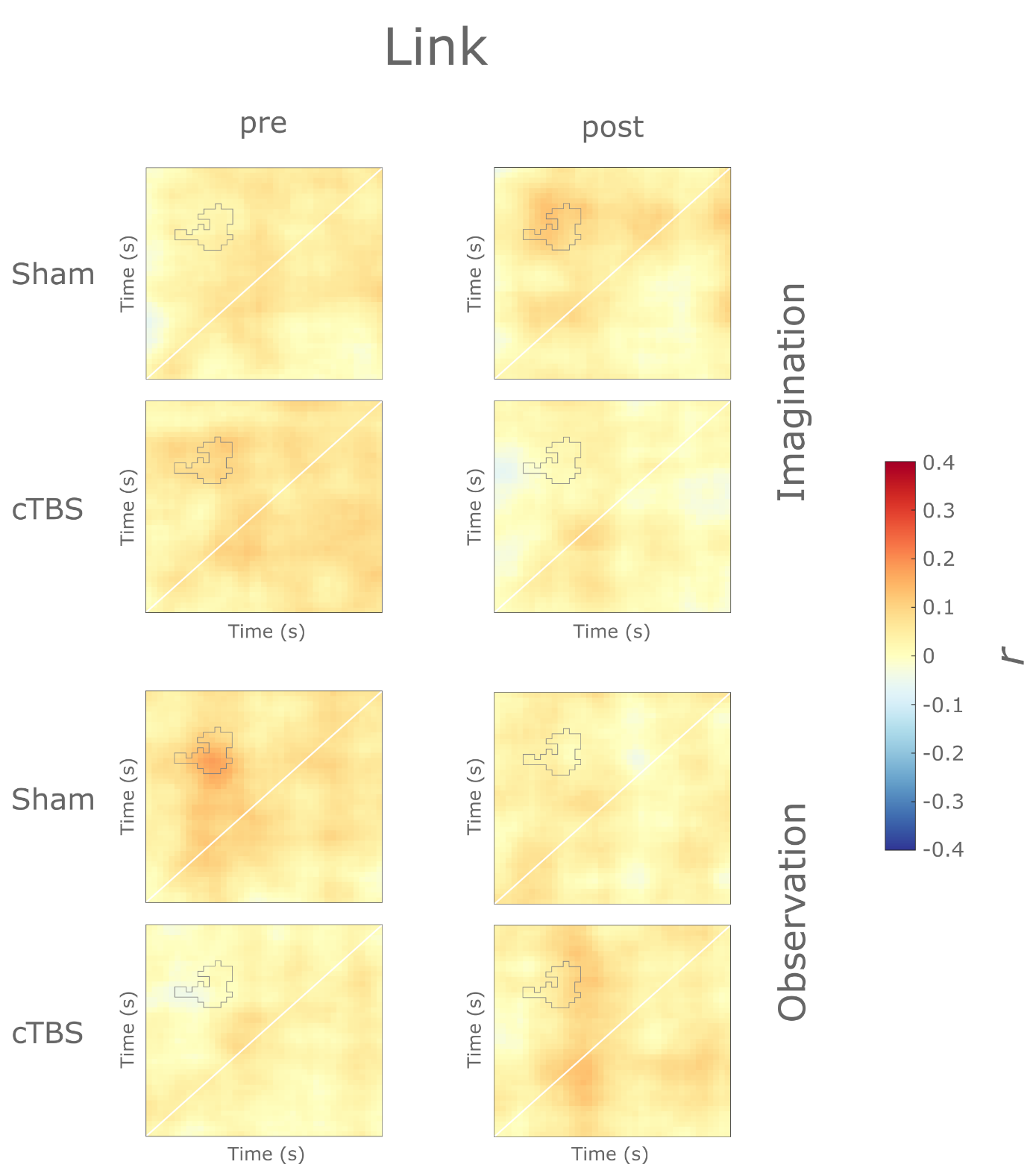

Figure 3 – figure supplement 1. Raw representational dissimilarity maps for linked events. Time × time similarity maps depict Pearson’s r between theta frequency vectors of event A and B at each time point. The grey line denotes the average significant cluster for the change from pre to post, the difference between imagination and observation and between the cTBS and the sham groups as found for linked events across the electrodes T7, TP7, and P7.

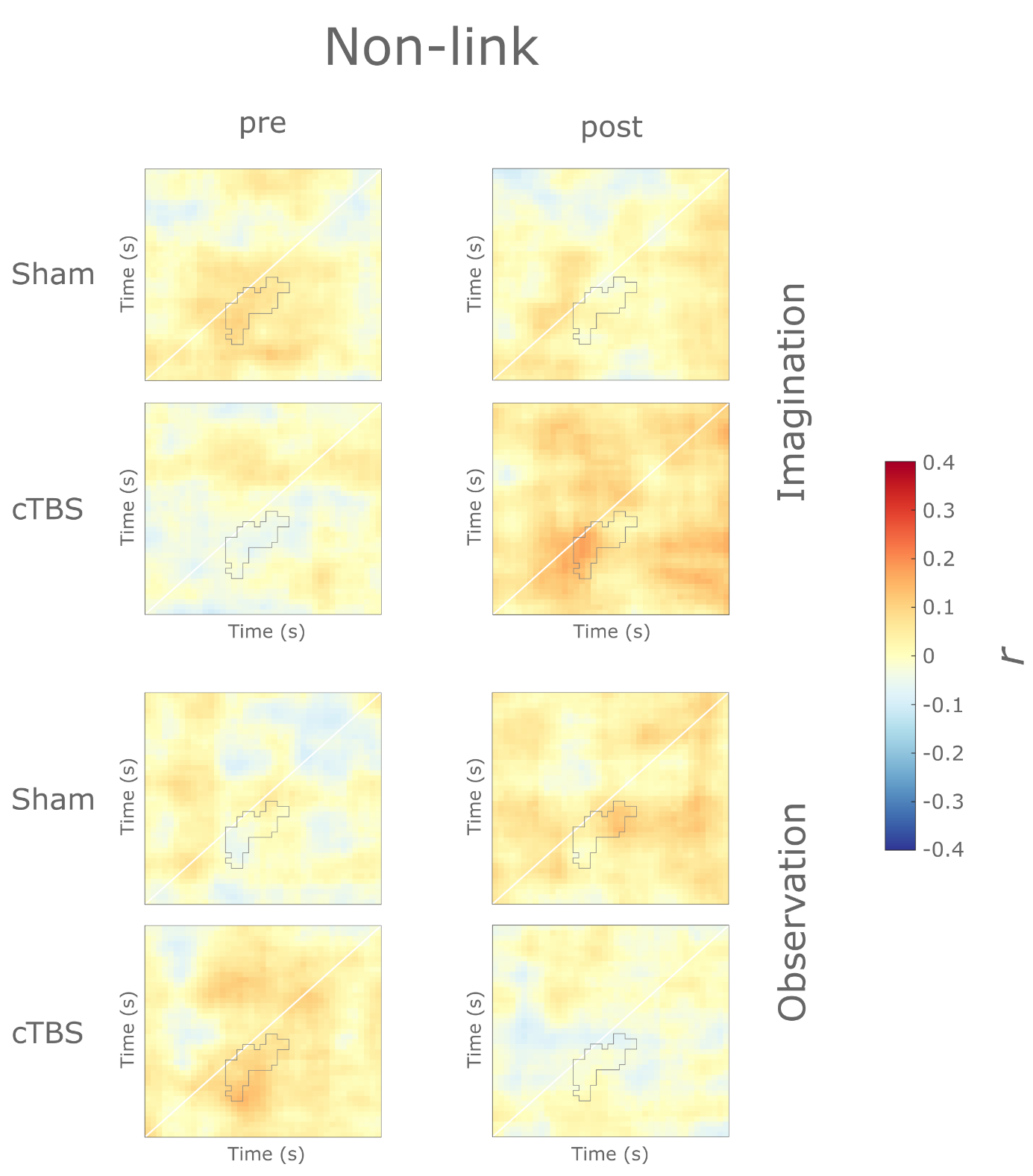

Figure 3 – figure supplement 2. Raw representational dissimilarity maps for non-linked events. Time × time similarity maps depict Pearson’s r between theta frequency vectors of event A and X at each time point. The grey line denotes the average significant cluster for the change from pre to post, the difference between imagination and observation and between the cTBS and the sham groups as found for non-linked events at the electrode FT7.
